# Disease-causing variant recommendation system for clinical genome interpretation with adjusted scores for artefactual variants

**DOI:** 10.1101/2022.10.12.511857

**Authors:** Ho Heon Kim, Junwoo Woo, Dong-Wook Kim, Jungsul Lee, Go Hun Seo, Hane Lee, Kyoungyeul Lee

## Abstract

**Background:** In the process of finding the causative variant of rare diseases (RD), accurate assessment and prioritization of genetic variants is essential. Although quality control (QC) of genetic variants is strictly performed, the presence of artefactual variants in the remaining set of variants can deteriorate the process. Variant QC and prioritization have been treated as separate processes, leading to limited efficiency and risk of misdiagnosis.

**Results:** We developed a disease-causing variant recommendation system that integrates quality control into variant prioritization by adjusting scores for artefactual variants. We confirmed that the QC-related features of the variants contribute to a significant performance improvement. For genomic data from 2,878 patients with rare disorders, the recall rate of finding causative variants was 0.961 for the top 5 ranked variants. We also found that our system recognized the anomaly of QC-related features, so that the scores of artifactual variants to be disease-causing were assessed relatively low.

**Conclusions:** Integration of variant QC and prioritization help reduce the risk of misdiagnosis based on artefactual variants and increase the effectiveness of clinical genome interpretation.

## Introduction

Rare disorders (RD), of which 80% have genetic causes, are estimated to affect approximately 6% of the global population [1, 2]. The advent of next-generation sequencing (NGS) has made a profound impact in the human genetics/genomics and medical genetics fields by revolutionizing the way rare disease diagnostics and disease gene discovery are performed [3]. The length of diagnostic odysseys of RD patients can be greatly shortened as genetic variants in all possible disease genes can be assessed simultaneously in an unbiased manner [4, 5]. However, finding the causal variant among hundreds of variants is a complicated task that requires gathering the most updated information from various public and private databases and assessing the variant’s pathogenicity in the context of a patient’s symptom similarity to the reported phenotypes of each gene/disease. Therefore, researchers have developed various computational tools to assist variant assessment for the effective diagnosis of RD patients [6].

One confounding factor affecting variant assessment is the presence of false variant calls or artefactual variant calls [7, 8]. Researchers have studied methods for removing those unexpected variants by performing quality control (QC) on genetic variants called from raw NGS data [9]. The QC process includes NGS read alignment/preprocessing, combining multiple variant calling tools, and rigorous filtering to remove false-positives [6]. Filtering methods, including GATK HaplotypeCaller and GATK UnifiedGenotyper [10], showed approximately 96% sensitivity and approximately 98% precision [11]. Additional processes will help better variant calling, such as base quality score recalibration (BQSR) and local realignment around indel variants [12]. Variant quality score recalibration (VQSR) of GATK is a classification method for QC training using a Gaussian mixture model [11]. However, it should be noted that many false variant calls remain even after QC and affect finding the causative variants [3]. Such false-positives can be assessed as top-ranked causative variants by variant prioritization, which can lead to an incorrect diagnosis based on artefactual variants [13]. Only a small fraction of false variants left in the data can considerably deteriorate the variant prioritization effectiveness.

Variant prioritization based on genotype™phenotype knowledge alongside variant data is typically used to find the causative variant(s) [3]. There are some tools, such as Exomiser [14], AMELIE [15], and LIRICAL [16], which use clinical phenotype data of patients in terms of Human Phenotype Ontology (HPO) to prioritize each candidate variant based on reference phenotypic knowledge. Additionally, variants can be filtered before prioritization so that the algorithm can assess only a reasonable number of variants. Exomiser filtered variants based on criteria such as the type of variants, allele frequencies, qualities of variants from the VCF file, a predefined set of genes, and inheritance patterns [14]. Although the variant filtering process helps improve variant prioritization accuracy, some diagnoses are filtered out because of low quality in the VCF, high allele frequencies, and incomplete penetrance [3].

Despite many attempts to address the problem of false calls affecting the diagnosis of RD patients, there is a fundamental limitation to previous approaches. Conventionally, variant QC and prioritization are considered two separate processes. As a result, a few remaining false-positives can be assessed as top-priority variants, and manual screening of variant artefacts for ranked variants is still needed. Not only does this decrease the efficiency of the diagnosis process using automated tools, but it also increases the risk of misdiagnosis based on artefact variants. Many false calls from NGS are scored incorrectly [17, 18], which are introduced by sequencing errors such as amplification bias during PCR [19] and polymerase mistakes. Nevertheless, if we apply strict quality standards to remove false-positives thoroughly, it is more likely that causative variants are also filtered out. Regardless of how accurate the following variant prioritization is, those removed variants cannot be assessed to diagnose the patient. Therefore, to avoid such a risk while reducing false calls among top-ranked variants predicted to be causative, we integrated features related to false-positives into a machine learning model for variant prioritization. In this study, we developed a recommendation system to find the disease-causing variants of RD patients, considering the risk of artefactual variants affecting patient diagnosis.

## Methods

### Study design and participants

In this retrospective study, we collected variant data generated in the process of clinical genetic interpretation from March 16, 2021, to Aug 3, 2022. Of the 15,526 suspected rare genetic diseases, we included 2,885 diagnosed patients. All known exonic regions of human genes (~22,000) were captured by xGen Exome Research Panel v2 (Integrated DNA Technologies, Coralville, Iowa, USA) and sequenced with Novaseq 6000 (Illumina, San Diego, CA, USA) as 150 bp paired-end reads. Exome sequencing data were aligned to the GRCh37/hg19 human reference genome using BWA-MEM, and variants were called by GATK v.3. after BQSR (base quality score recalibration) and local realignment with GATK (v.3.8) to reduce false-positive variant calls [20]. Variants were then annotated with Ensembl Variant Effect Predictor (VEP) and classified according to the American College of Medical Genetics and Genomics (ACMG) guidelines (Richards et al., Genet Med. 2015 May). The sequencing data were analysed using 3billion’s bioinformatics pipeline ‘EVIDENCE’ as previously described [5]. In addition, this study adhered to the TRIPOD statement on reporting predictive models [21].

### Development of rare disease causative variant recommendation system

We developed a recommendation system that feeds multivariate as follows: posterior probability for pathogenicity based on ACMG/AMP variant classification, disease semantic similarity between patient symptoms and disease phenotypes, and statistics related to QC (quality control) including VAF (variant allele frequency), DP (depth), QUAL (mapping quality), MQ (mapping quality), and variant allele frequency in the inhouse dataset. For a detailed description of feature engineering, first, the posterior probability was calculated using rules derived from EVIDENCE with 0.1 of prior probability and 350 odds of pathogenicity for the “very strong” category [22]. Second, disease semantic similarity refers to the ontology similarity between patient symptoms and rare disease phenotypes based on Human Phenotype Ontology (HPO, v.1.2). Disease semantic similarity was calculated as the average depth of their most informative common ancestor (MICA) between two trees on both sides (disease to symptoms; symptoms to disease) [5].

The model was built with random forest (RF), which is a machine learning algorithm based on ensembles of trees (weak classifiers) using bagging [23]. In total, 3,264 and 184,110,909 confirmed and unconfirmed causative variants with disease were collected, respectively. To overcome the class imbalance that confirmed causative variants is one or two variants among many variants in VCF, the model was trained with adjusted class-weight in sklearn 1.1.2.

### Model performance evaluation

To measure the performance of the recommendation system, we conducted the following experiments: 1) a conventional performance examination based on the collected dataset by splitting the training and test data, 2) a comparison of performance between other prioritizing models, including LIRICAL and Exomiser, and 3) an ablation test to determine the contribution of QC-related statistics to model performance.

To define the model performance, the hit rate (recall), which is a conventional metric for recommendation systems, was evaluated. It is referred to as the percentage of patients with at least one correct recommendation up to the cut-off k-rank, denoted as R@k [24]. For example, if a system predicted a variant for every 7 patients within 5-rank among 10 patients, R@5 is 0.7. Because some patients with dual diagnoses have multiple confirmed causative variants, model performance is measured in two ways. One method considers the case where at least one causal variant is predicted within the k-rank as a hit (from now on, we note this “single match”), and the other allows all the causal variant of patients to be predicted within it the k-rank as a hit (from now on, we note this “full match”).

First, to evaluate the consistency in the model’s performance, 5-fold cross validation (training set: 80%, test set: 20%) was performed. Because it is also essential to recognize concordance between the symptoms of patients and disease phenotypes in RD diagnosis practice, we evaluated the recall of models when the model predicted the pairs of variants and diseases correctly [25, 26].

Second, we compared the recall of each model based on the level of genes from the LIRICAL and Exomiser results, although our model can prioritize single variants instead of genes. To evaluate the performance of the other two models, we defined the probability of a gene as the highest probability of a variants of the gene (variant-level comparison included in supplementary 1).

Finally, we conducted an ablation test to identify the contributions of QC-related statistics to model performance. We compared two models: one was based on the model trained from all features, and the other was the model trained from features excluding QC-related statistics as features. Then, to measure the performance improvement, we conducted the Mann□Whitney U test, a popular nonparametric alternative for a t test, considering the recall value obtained from each model for the 5-fold dataset as observations of two groups [27]. In this process, we compared the best model performance with and without QC-related features among various machine learning models, including fully connected networks, random forest, and logistic regression.

### Model interpretation

For reliable AI, we conducted post hoc analysis with model-agnostic XAI (eXplainable AI) techniques, including SHAP (Shapley additive explanations) and permutation feature importance. SHAP provides an importance value (SHAP value) of each feature for a particular prediction by estimating the conditional expectation of features [28]. In addition, permutation feature importance, as known as the mean decrease in accuracy (MDA), refers to the magnitude of the average decrease in accuracy by shuffling values in a column.

## Results

### Demographic characteristics

The demographic characteristics of the patients are shown in Table 1. Of 2,878 patients, 1,605 were male in our retrospective cohort (55.6%). In addition, the proportion of infant patients was less than 50%. Most of their symptoms were in the nervous system (n=1,268, 44.0%), followed by the musculoskeletal system (n=962, 33.3%).

**Table 1.** Demographic and clinical characteristics of the patient cohort

| Variables, |  | Patients with<br>confirmed causative variants<br>(n=2,878) |
| --- | --- | --- |
| Gender, (n, (%)) | Male | 1605 (55.6) |
|  | Female | 1280 (44.4) |
| Onset age, (n, (%)) | Infancy | 804 (27.9) |
|  | Neonatal | 676 (23.4) |
|  | Childhood | 607 (21.0) |
|  | Adult | 532 (18.4) |
|  | Antenatal | 165 (5.7) |
|  | Adolescent | 87 (3.0) |
|  | Unknown | 7 (0.2) |
|  | Infant | 4 (0.1) |
|  | Elderly | 3 (0.1) |
| Symptoms | Nervous system | 1268 (44.0) |
|  | Musculoskeletal system | 962 (33.3) |
|  | Head or neck | 704 (24.4) |
|  | Cardiovascular system | 678 (23.5) |
|  | Eye | 643 (22.3) |
|  | Integument | 512 (17.7) |
|  | Growth | 485 (16.8) |
|  | Metabolism/homeostasis | 429 (14.9) |
|  | Limbs | 399 (13.8) |
|  | Genitourinary system | 379 (13.1) |
|  | Ear | 357 (12.4) |
|  | Digestive system | 327 (11.3) |
|  | Endocrine system | 229 (7.9) |
|  | Immune system | 219 (7.6) |
|  | Blood and blood-forming tissues | 202 (7.0) |
|  | Respiratory system | 117 (4.1) |
|  | Neoplasm | 99 (3.4) |
|  | Prenatal development or birth | 54 (1.9) |
|  | Constitutional symptom | 38 (1.3) |
|  | Breast | 25 (0.9) |
|  | Cellular | 20 (0.7) |
|  | Voice | 13 (0.5) |

### Model performance

In 5-fold cross validation, the model showed an average recall of 0.712 and 0.624 at Rank-1 for single matches and full matches to confirmed causative variants, respectively. At 5-rank, an average of 0.961 and 0.921 of recall were shown in both cases. Over 5 k, the recall score is saturated (Figure 1-A). Among our proposed model, LIRICAL, and Exomiser, our proposed model showed superior performance at any k, followed by LIRICAL and Exomiser. Our model showed 0.876 and 0.957 recalls at 1 and 3 k, respectively (Figure 1-B).

**Figure 1.**
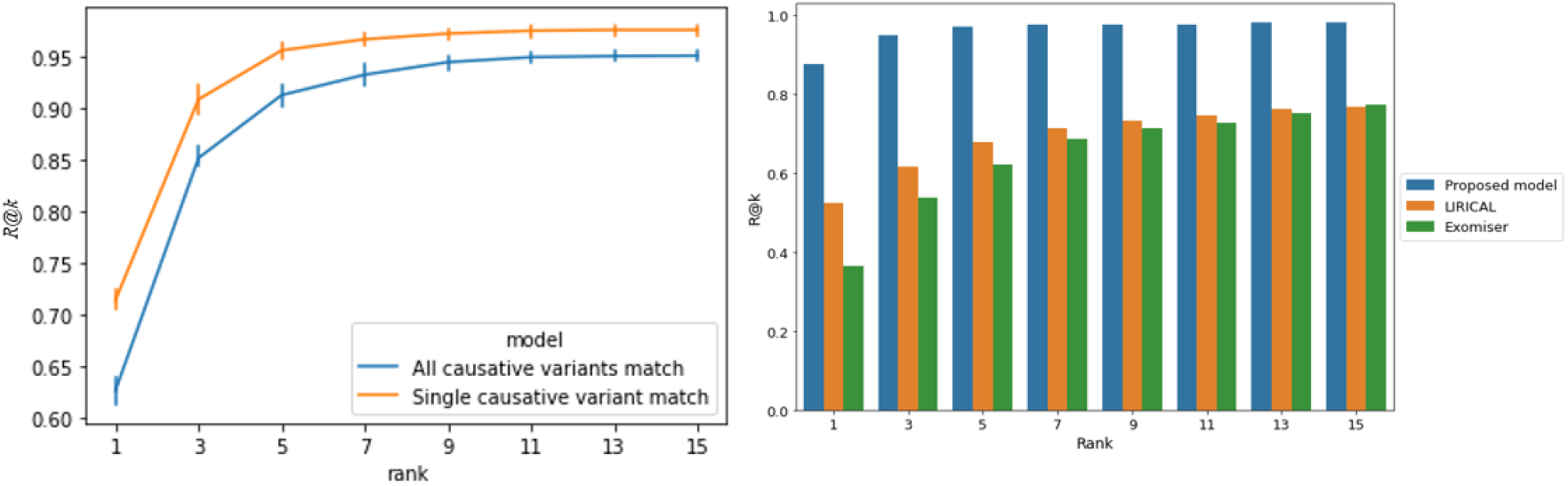
A) Model performance based on single match (single causative variant) and full match (all causative variants match) B) Comparison of recall of Exomiser, LIRICAL, and proposed model by gene-level match.

### Ablation test

In an ablation test to identify the contribution of statistics related to QC as features, recall at 1-rank was 0.699 and 0.480 in models trained with and without QC statistics, respectively (Figure 2-A). Model-trained data with QC statistics showed significantly superior performance than the model without those on both single matches and full matches (p<0.05, at all k values) (Figure 2-A, B).

**Figure 2.**
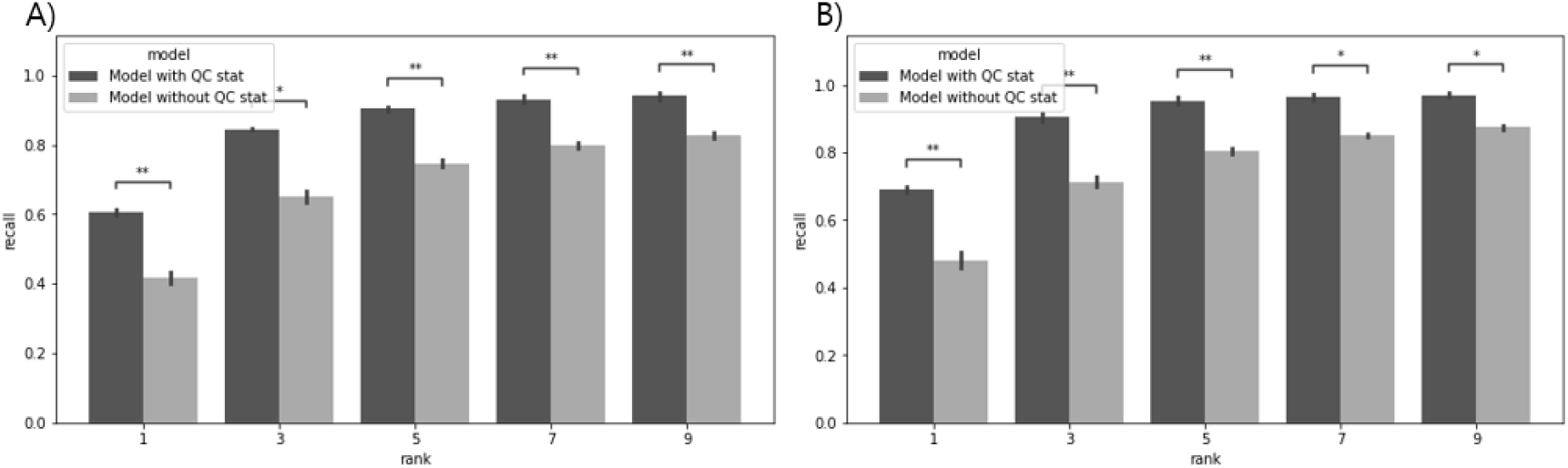
Ablation test of QC statistics contribution for model performance.

### Post hoc interpretation

Figure 3 shows SHAP summary plots for the model’s features. A relatively high value of ACMG posterior, disease semantic similarity and VAF in the dataset showed a positive SHAP value that contributed to the model predicting a high probability for causative confirmed variants with disease. In contrast, a low value of inhouse allele frequency was estimated as a positive SHAP value (Figure 3-A). If other variables were constant, shuffling the ACMG posterior probability caused the greatest decrease in accuracy, followed by disease semantic similarity (Figure 3-B).

**Figure 3.**
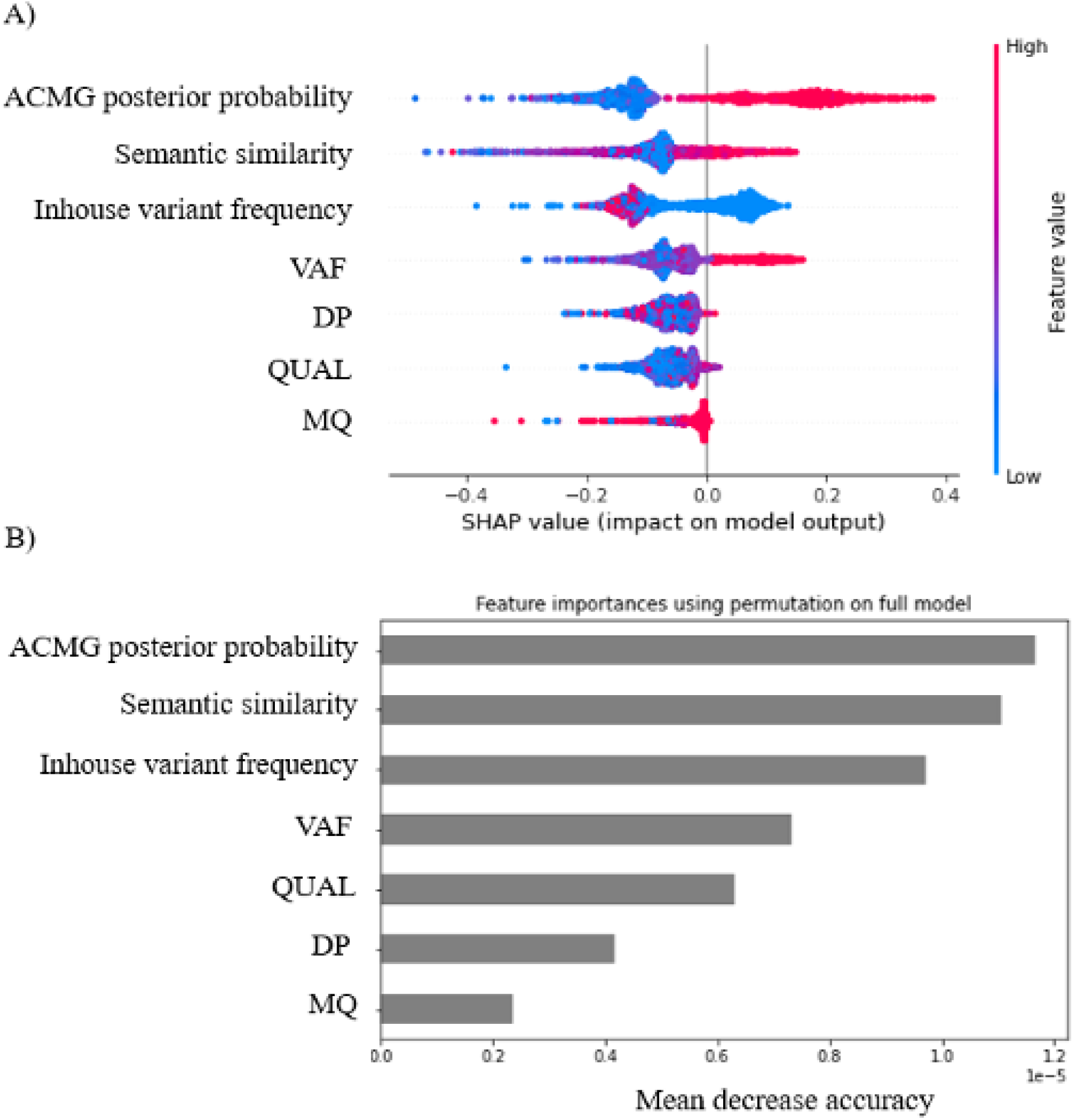
SHAP summary plots for the feature importance (A) and bar plot of mean decrease in accuracy (MDA) for each feature (B).

### Individual case

For each model, we compared the predicted rank of confirmed causative variants in the test dataset. The model trained with variant QC-related statistics predicted the rank of causative variants in the top 5 rank than the model without these statistics. Also, in comparison between our model and other models (LIRICAL, and Exomiser), our model predicted most of the confirmed variants within 5th rank (Figure 4-A).

**Figure 4.**
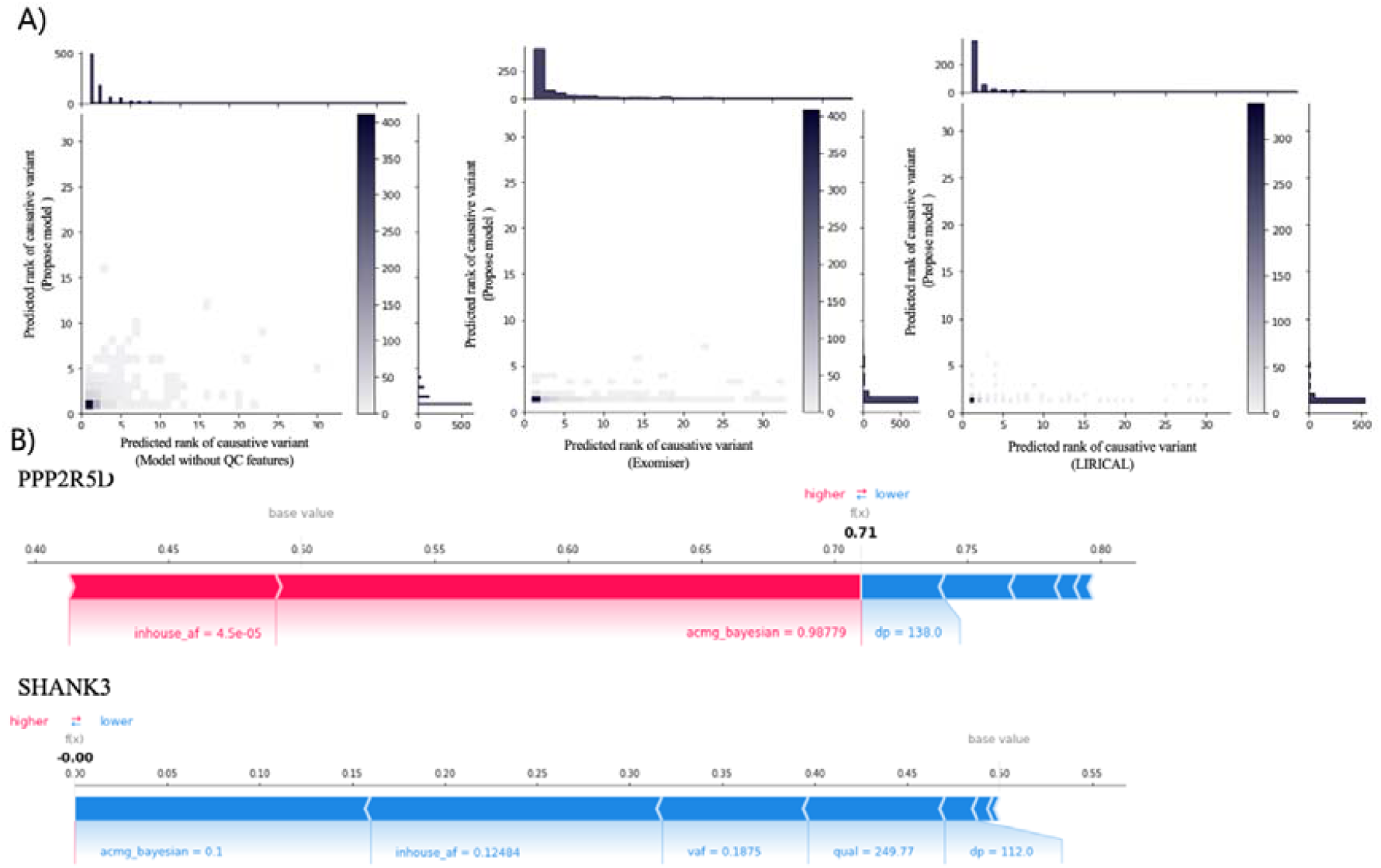
A) Distribution of predicted rank of causative variant from each model. B) SHAP plot for the contribution of features in our model. (A) the model with QC-related statistics predicted most of causative variants in top 5 compared to three models (model without QC-related statistics, Exomiser, LIRICAL) in the test dataset. The histogram of top and right of center plot showed the number of patients with predicted rank of causative variant (B) Features of variants in the PPP2R5D gene; those of false call variants in the SHANK gene. The model predicted a 0.71 probability of PPP2R5D for confirmed causative genes. From 0.5, a low level of inhouse allele frequency (0.00045) and a high score of posterior pathogenic probability based on ACMG (0.98779) contributed to increasing the probability as the length of red bars. For SHANK3, which was represented by false call variants, QC-related features contributed to decreasing probability as each blue bar.

For individual case, a patient with seizure and autistic behaviour had NM_006245.4:c.612G>C and was diagnosed with an intellectual developmental disorder caused by in PPP2R5D. By aggregating variants into genes, our model predicted this variant as the highest rank at 1 with an 0.71 probability for disease-causing genes. our model predicted the probability SHANK3 as 0, which contains the false call variant was predicted. However, LIRICAL predicted this gene at the 8th rank with 0.0017 of probability, and SHANK3 was found in the 5th rank with 0.022 of probability (Figure 4-B).

## Discussion

Some limitations in this study should be improved upon in the future. First, all the patient data used in this study are based on inhouse genome data from an RD diagnosis corporation, 3billion. Therefore, there can be possible overfitting of the model to the NGS sequencing and diagnosis process of 3billion. Although around 4,000 diagnosed cases from UK 100,000 Genome Project could have been used to evaluate the model, the data are not publicly available [29]. We expect that this model will be evaluated without bias as different sources of patient genome data are published in the future. In addition, we selected the features related to false calls, such as allele frequency, QUAL, and DP, which are also widely used for QC of the variants; therefore, the model cannot account for other underlying risk factors introducing high scores on the artefactual variants. There may be heterogeneous risk factors affecting the prioritization of artefacts depending on the target gene, such as homopolymer regions or pseudogenes. The model may be further optimized by applying state-of-the-art methods of machine learning and artificial intelligence such as sequence-based input, multitask learning, or semisupervised learning. Nevertheless, we expect that the findings in this study and the trained model will help other researchers and clinicians find the causative variant(s) more effectively for their patients.

We developed the recommendation system for disease-causing variants with the most relevant disease. Our system showed superior performance to other prioritization models by leveraging posterior probability for pathogenicity based on ACMG guidelines, disease semantic similarity, and variant calling QC-related statistics. As a result, our system lowered the probability of false call variants or genes causing disease to be low. By integrating both prioritization and false call adjusting processes, our system evaluated the rank of causative variants as high and lowered that of false call variants, unlike other false call-filtering models.

As a data-driven approach, our system has shown high performance in predicting confirmed causative variants by leveraging data from a variant calling pipeline without a false call variant filtering process. Rather than focusing on excluding false call variants, our system used the QC statistics as covariates to predict causative variants accurately. Figures 2 and 3 imply that our system learned the pattern that causative variants with high allele frequency in the inhouse dataset, homopolymer variants, and low VAF were less likely to be confirmed. For an individual case, we confirmed this mechanism of our system by SHAP value indirectly (Figure 4).

Naturally, in NGS covering a large number of base pairs, the rule-based filtering methodology for false call variants has a limitation. Although no matter how sophisticated rules are set, false call variants could remain, and unexpected filtering of true call variants may occur. Accordingly, hard filtering with rule-based approaches excluding low-quality variants and unusually high minor allele frequencies cannot address filtering out false call variants entirely [9]. Even in our experiment, some confirmed causative variants were ruled out, which could deprive the opportunity of clinical genetics to further explore variants such as Sanger sequencing. Therefore, our data-driven approaches with the adjusted ranks for false call variants can improve clinical genomic interpretation, providing the opportunity to assess all candidates of variants with high accuracy.

We additionally conducted a comparison between false call variants and all of the variants in randomly chosen samples (Supplementary 2). Some false call variants were manually annotated under criteria including a low level of VAF and the presence of a low complexity region, homopolymer variant, and high AF in the inhouse dataset by clinical geneticists. The true call variants were defined as the complementary set of false call variants and can imply that other false call variants can be included.

We also found many false call variants in the clinical interpretation process at the endpoint of genome analysis, even if our NGS analysis pipeline adheres to best practices. Because two distributions have an overlapping region, true call variants were excluded unexpectedly despite the adoption of a sophisticated cut-off. Instead, our system predicted a lower probability of false calls by leveraging QC statistics as covariates. Therefore, we can expect a ceteris paribus effect in our model, which is that our system will score true call variants with a high priority when the ACMG posterior probability and disease semantic similarity of a true call variant were equal to false call variants [30].

## List of abbreviations

RD: Rare disorders
NGS: next-generation sequencing
QC: Quality control

## Declarations

### Ethics approval and consent to participate

All the data in this study had the consents from the participants that it can be used for the research purpose.

### Consent for publication

All the data in this study had the consents from the participants that it can be used for the research purpose.

## Availability of data and materials

The data and codes used to train and evaluate the model will be uploaded in the repository.

Project name: Resys4RD

Project home page: https://github.com/4pygmalion/Resys4RD

Programming language: Python

License: MIT

## Authors’ contributions

HHK designed the study, implemented the methods, performed the experiments and data analysis, and draft the manuscript. JW and DWK implemented the methods and draft the manuscript. JL, GHS, and HL designed the study and participated in writing the manuscript. KL participated in the study design and data analysis, and draft the manuscript.

