## Supplementary Figures for "Disease-causing variant recommendation system for clinical genome interpretation with adjusted scores for artefactual variants"

- Mailing address: 14th floor, 416, Teheran-ro, Gangnam-gu, Seoul, 06193 Seoul, Republic of Korea

- e-mail address:

Ho Heon Kim <>, Junwoo Woo <>, Dong-Wook Kim <>, Jungsul Lee < >, Go Hun Seo <>, Hane Lee <>, Kyoungyeul Lee <>

### Supplementary 1

We attach the performance comparison of three models, including our proposed model, LIRICAL, and Exomiser, based on the variant level. For our dataset, LIRICAL and Exomiser showed extremely low recall compared to our model. This may be because LIRICAL provides the pathogenic probability of a variant to one decimal point, or Exomiser may internally filter out many variants.

Supplementary figure 1. Comparison of performance at the variant level among the proposed model, Exomiser, and LIRICAL

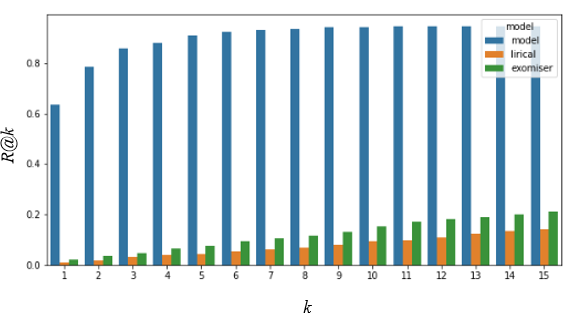

### Supplementary 2

We additionally conducted a comparison between false call variants and all of the variants in randomly chosen samples (Supplementary 2). Some false call variants were manually annotated under criteria including a low level of VAF and the presence of a low complexity region, homopolymer variant, and high AF in the inhouse dataset by clinical geneticists. The true call variants were defined as the complementary set of false call variants and can imply that other false call variants can be included.

Supplementary figure 2. The distribution of DP and VAF between false calls and true calls (soft-labelled)

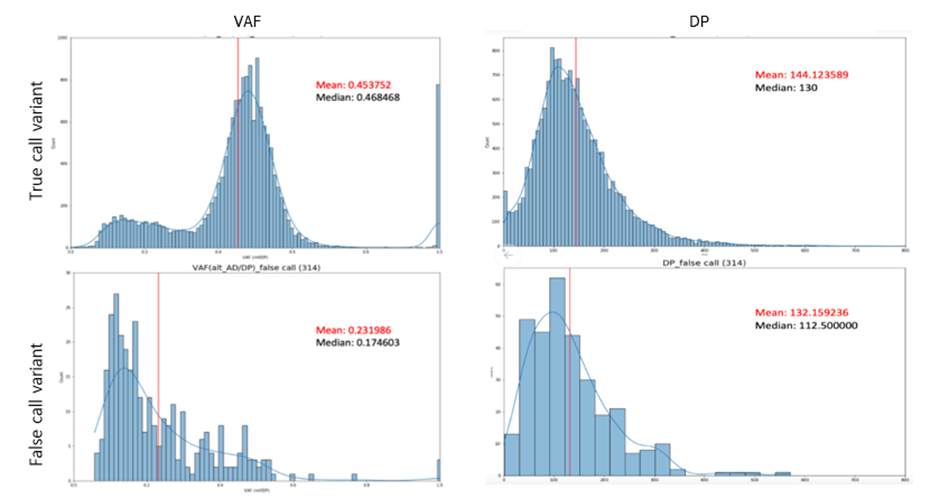
